# Ecological diversity of *Anopheles gambiae s.l.* and insecticide resistance across refugee camps in Kenya

**DOI:** 10.1101/2025.10.13.682237

**Authors:** Martin K. Rono, Caroline Wanjiku, Brian Bartilol, Audrey Oronda, Adam Nduni, Garama Kenga, Mercy Tuwei, Lynette I. Ochola-Oyier, Robert W. Snow, Joseph Mwangangi, Marta Maia

## Abstract

Malaria remains a major threat during humanitarian crises, necessitating targeted vector control strategies informed by local vector dynamics. Between May and July 2023, we conducted larval surveys in refugee settlements across Dadaab, Kakuma, and Kalobeyei (Kenya), collecting Anopheles larvae. Genotyping of 728 specimens revealed spatial variations in species composition. Overall, *Anopheles arabiensis* was the dominant species (59%, n=426), followed by *Anopheles coluzzii* (35%, n=252), and *Anopheles rufipes* (1%, n=7). In Dadaab, *An. arabiensis* was overwhelmingly dominant (94%, n=352/374). In contrast, the Kakuma/Kalobeyei complex was characterized by the co-occurrence of *An. coluzzii* (72%, n=252/350) and *An. arabiensis* (22%, n=74/350), with *An. rufipes* exclusively found in Kalobeyei (7%, n=6/89) (Figure 2B). Notably, no members of the *Anopheles funestus* group or *Anopheles stephensi* were detected. However, approximately 5% of the larvae across the sites could not be resolved molecularly. High frequencies of the *L1014F kdr* mutation, a pyrethroid resistance marker, were detected in *An. coluzzii* (Kakuma: 50%; Kalobeyei: 63%) and *An. arabiensis* (Kakuma: 10%; Kalobeyei: 30%) populations in Turkana County. Interestingly, no *kdr* mutations were observed in the *An. arabiensis* population from Dadaab. These findings highlight significant spatial diversity in vector species composition and resistance profiles, with *An. coluzzii* emerging as a dominant, pyrethroid-resistant vector in the Kakuma/Kalobeyei complex. The results underscore the urgent need for targeted interventions, including resistance monitoring and alternative insecticide-based strategies, to mitigate malaria transmission risks in fragile, humanitarian settings. Further studies are warranted to address unidentified larval species and seasonal transmission dynamics.

## Introduction

Conflict and the displacement of large populations are recognized as significant public health risks. The rapid influx of refugees, coupled with limited sanitation infrastructure, often leads to disease outbreaks. Infectious diseases are a major cause of morbidity and mortality in refugee camps. Between 2009 and 2017, the United Nations High Commissioner for Refugees (UNHCR) reported over 350 infectious disease outbreaks in refugee camps globally[1, 2], with Kenya, Chad, and Thailand among the countries bearing the highest disease burdens. Vector-borne diseases, including yellow fever, dengue, and malaria, contribute substantially to this burden during humanitarian crises[3–6].

In Kenya, Kakuma and Dadaab refugee camps were established in response to humanitarian crises stemming from conflict and famine in neighboring countries. Kakuma, located in Turkana County in northwestern Kenya, was established in 1992 and is home to approximately 300,000 inhabitants, primarily refugees from South Sudan and Somalia[7, 8]. In 2016, the UNHCR, in collaboration with the national and county governments of Turkana, established the Kalobeyei Integrated Settlement to alleviate congestion in Kakuma and promote socio-economic integration with the host community.

Dadaab, established in 1991, is the largest refugee camp in Kenya, located in Garissa County near the Kenyan-Somali border. With a population of approximately 400,000, predominantly Somali refugees, Dadaab is administratively divided into three sub-camps: Ifo, Hagadera, and Dagahaley[9]. These camps face significant challenges in managing public health risks, particularly vector-borne diseases, due to overcrowding, limited resources, and the dynamic nature of refugee populations.

Turkana County, Kenya, despite its semi-arid climate and low rainfall, experiences a disproportionately high malaria burden[10–12], sustained by local, year-round transmission that shows little correlation with rainfall[13]. This persistent threat is now compounded by the recent discovery of two novel malaria vectors uniquely adapted to arid ecologies: the invasive *Anopheles stephensi* [14]and the West African *Anopheles coluzzii* [15]. The larval ecology of *An. stephensi*, in particular, is characterized by its specialization for breeding in a wide range of human-made water containers including storage tanks, cisterns, and discarded tires which are abundant in urban settings and essential for water security in drought-prone areas.. This adaptability is further enhanced by its independence from rainfall and its ecological flexibility to utilize natural sites like riverbeds. Together, these traits underpin its role as a major and emerging urban malaria vector in Africa [16–18]. Similarly, *An. coluzzii* is strongly associated with anthropogenic habitats in arid ecologies, with larvae typically found in permanent or semi-permanent human-made aquatic habitats such as irrigated fields, urban drainage, and stored water containers[19]. These adaptations make refugee camps like Kakuma particularly vulnerable. Essential water harvesting and storage practices create abundant breeding sites, a factor strongly linked to increased mosquito-borne disease transmission[20–22].The coexistence of *An. coluzzii* and *An. stephensi* with established vectors like *An. arabiensis* suggests a complex and evolving transmission landscape, potentially exacerbating malaria burden in Turkana and neighboring regions.

Understanding the bionomics and insecticide resistance profiles of these vectors is critical for refining disease control strategies and especially within humanitarian contexts. In this study, we examined the diversity and insecticide resistance profile of local malaria vector populations of Kakuma and Dadaab refugee camps.

## Methods

### Larval surveys

Mosquito larvae were collected from randomly selected sites in Kakuma and Dadaab refugee camps between May 2023 and July 2023 (Supplementary Table 1). Habitats were sampled once during the study period. Larvae at the L1-L3 developmental stages were sampled from a variety of habitats, including water containers, stagnant water, and roadside pools, using a standard 350 mL dipper. At each site, at least three dips were taken to quantify larval density. Anopheles larvae were separated and individually preserved in 1.5 mL microcentrifuge tubes containing 95% ethanol for transport to the KEMRI Wellcome Trust Research Programme in Kilifi, Kenya.

### Nucleic acid extraction and molecular analysis for species identification

Field-collected larvae were rinsed with nuclease-free water, and genomic DNA extracted using the Chelex method. Briefly, whole larvae were transferred into individual 1.5 mL microcentrifuge tubes containing 50 µL of 20% Chelex resin (Bio-Rad, USA) and homogenized using polypropylene pestles. The lysate was incubated at 100 °C while shaking at 650 rpm on a ThermoMixer (Eppendorf, Hamburg, Germany). The solution was centrifuged at 10,000 × *g* for 2 minutes, and the supernatant transferred to a new 1.5 mL microcentrifuge tube. This process was repeated twice, and the extracted DNA stored at −80 °C until further analysis. *Anopheles gambiae s*.*l*. sibling species were identified using a previously described PCR method targeting the intergenic spacer (IGS) region of the ribosomal DNA [23]. Samples that did not amplify for *An. gambiae s*.*l*. were screened for members of the *An. funestus* complex using established protocols[24] and *An. stephensi* using established protocols [14].

### Nucleic acid extraction and molecular analysis for species identification

Field-collected larvae were rinsed with nuclease-free water, and genomic DNA was extracted using the Chelex method. Briefly, whole larvae were transferred into individual 1.5 mL microcentrifuge tubes containing 50 µL of 20% Chelex resin (Bio-Rad, USA) and homogenized using polypropylene pestles. The lysate was incubated at 100 °C while shaking at 650 rpm on a ThermoMixer (Eppendorf, Hamburg, Germany). The solution was centrifuged at 10,000 × ^*^g^*^ for 2 minutes, and the supernatant was transferred to a new 1.5 mL microcentrifuge tube. This process was repeated twice, and the extracted DNA was stored at −80 °C until further analysis.

Species identification was performed using a sequential molecular workflow. First, *Anopheles gambiae s*.*l*. sibling species were identified using a PCR method targeting the intergenic spacer (IGS) region of the ribosomal DNA [23]. Samples identified as *An. gambiae s*.*s*. were further characterized to distinguish between the M (*An. coluzzii*) and S (*An. gambiae*) forms via a PCR assay targeting the SINE200 insertion/deletion polymorphism on the X chromosome [25]. The primers used were: Forward: 5′-GTGTGCACCTCGACGTAC-3′ and Reverse: 5′-CGGAGTGACCAGGACACC-3′. PCR products were resolved on a 2% agarose gel; samples showing a band at ~315 bp were classified as *An. coluzzii*, those with a band at ~250 bp as *An. gambiae*, and those with both bands as hybrid forms. Samples that did not amplify for *An. gambiae s*.*l*. were subsequently screened for members of the *An. funestus* complex [24] and for *An. stephensi* [14]. This sequential identification workflow was chosen based on historical data and previous studies indicating the dominance of the *An. gambiae* complex over the *An. funestus* group in these arid ecological zones [26].

### Molecular Identification of PCR-Unresolved Mosquito Specimens

For mosquitoes unresolved by initial PCR, the ITS2 region was amplified using primers from Beebe & Saul (1995)[2]. The 10 µL PCR reaction contained 5 µL of 2X GoTaq® Master Mix (Promega, USA), 0.5 µL of each primer, 1.5 µL of DNA, and 2.5 µL nuclease-free water.

Cycling conditions were: 95°C for 5 min; 40 cycles of 95°C for 15 sec, 52°C for 20 sec, and 72°C for 1 min; with a final 72°C for 10 min. Amplicons were purified with the QIAquick PCR Purification Kit (Qiagen, Germany). Sanger sequencing used BigDye Terminator v3.1 (Applied Biosystems, UK) on an ABI 3730xl sequencer. Chromatograms were edited in CLC Main Workbench 24. Consensus sequences were identified via BLASTn against the NCBI nt database under default parameters[27].

### Insecticide resistance profiling

The presence of point mutations at position 1014 of the voltage-gated sodium channel (vgsc) gene (knock down resistance (*kdr*) mutations) were investigated using a TaqMan probe-based quantitative real-time PCR (qPCR) assay. The assay utilized one set of primers: Forward: 5′-CAT TTT TCT TGG CCA CTG TAG TGA T-3′, Reverse: 5′-CGA TCT TGG TCC ATG TTA ATT TGC A-3′ and three probes for detection of: wild-type allele: 5′-CTT ACG ACT AAA TTT C-3′ (labeled with HEX fluorophore), Vgsc-L1014F mutation: 5′-ACG ACA AAA TTT C-3′ (labeled with FAM fluorophore) and Vgsc-L1014S mutation: 5′-ACG ACT GAA TTT C-3′ (labeled with FAM fluorophore). The qPCR cycling conditions consisted of an initial denaturation at 95 °C for 10 minutes, followed by 40 cycles of denaturation at 95 °C for 10 seconds and annealing and extension at 65 °C for 45 seconds. A sample is considered positive for a specific allele (wild-type, L1014F, or L1014S) if its associated probe generates a fluorescence signal above the threshold during qPCR.

## Results

### Vector abundance, distribution and taxonomic assignment

Larval surveillance conducted in May and July 2023 identified productive *Anopheles* breeding habitats in both the Dadaab (Garissa County) and Kakuma/Kalobeyei (Turkana County) refugee camps (Figure 1A). The survey of 13 habitats revealed an overall mean density of 6.6 larvae/dip, though this varied significantly between camp environments (Table 1).

**Figure 1:**
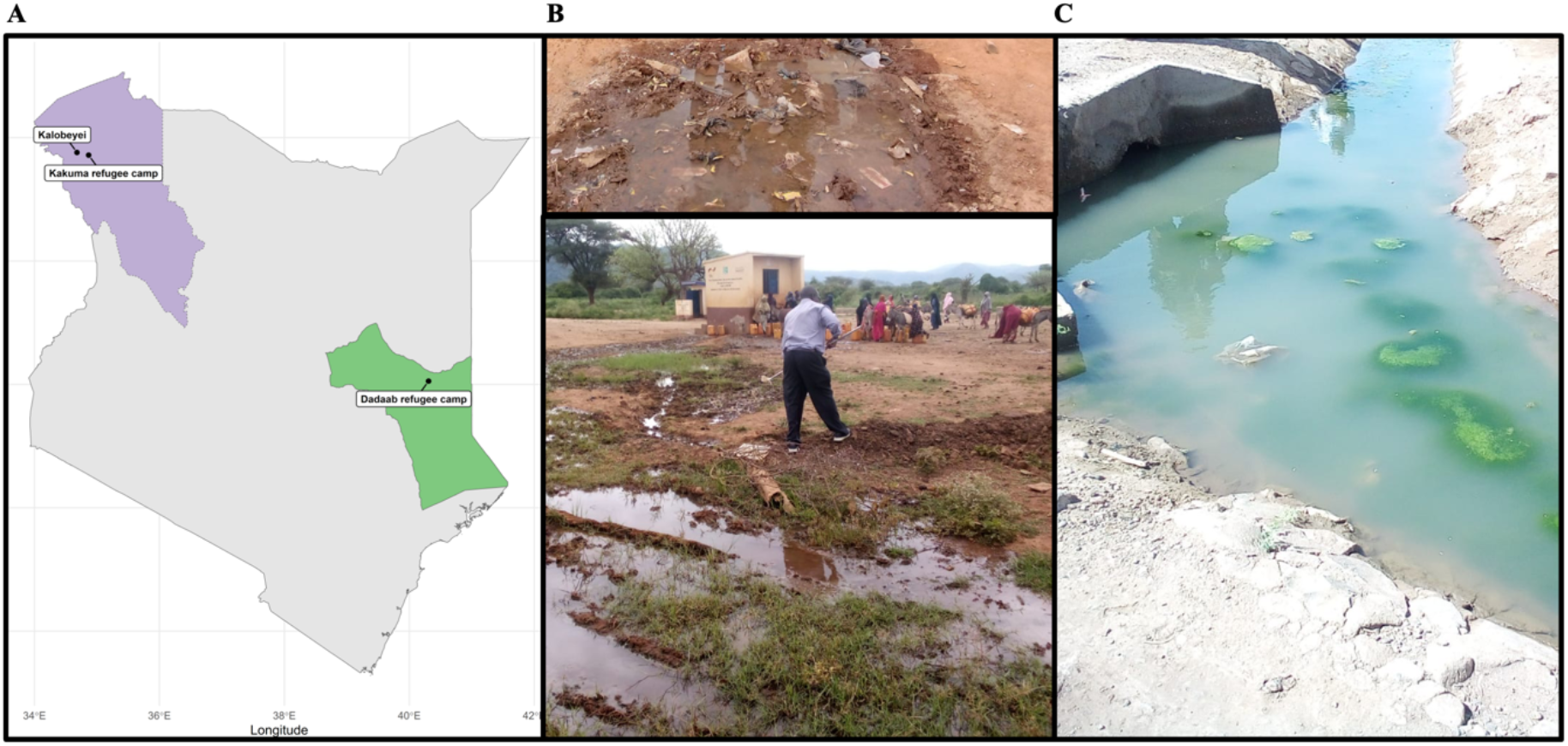
Malaria vector larval sampling in Kakuma and Dadaab refugee complex. A)Map of sites sampled for anopheline larvae; B and C) larval breeding sites

**Table 1.**
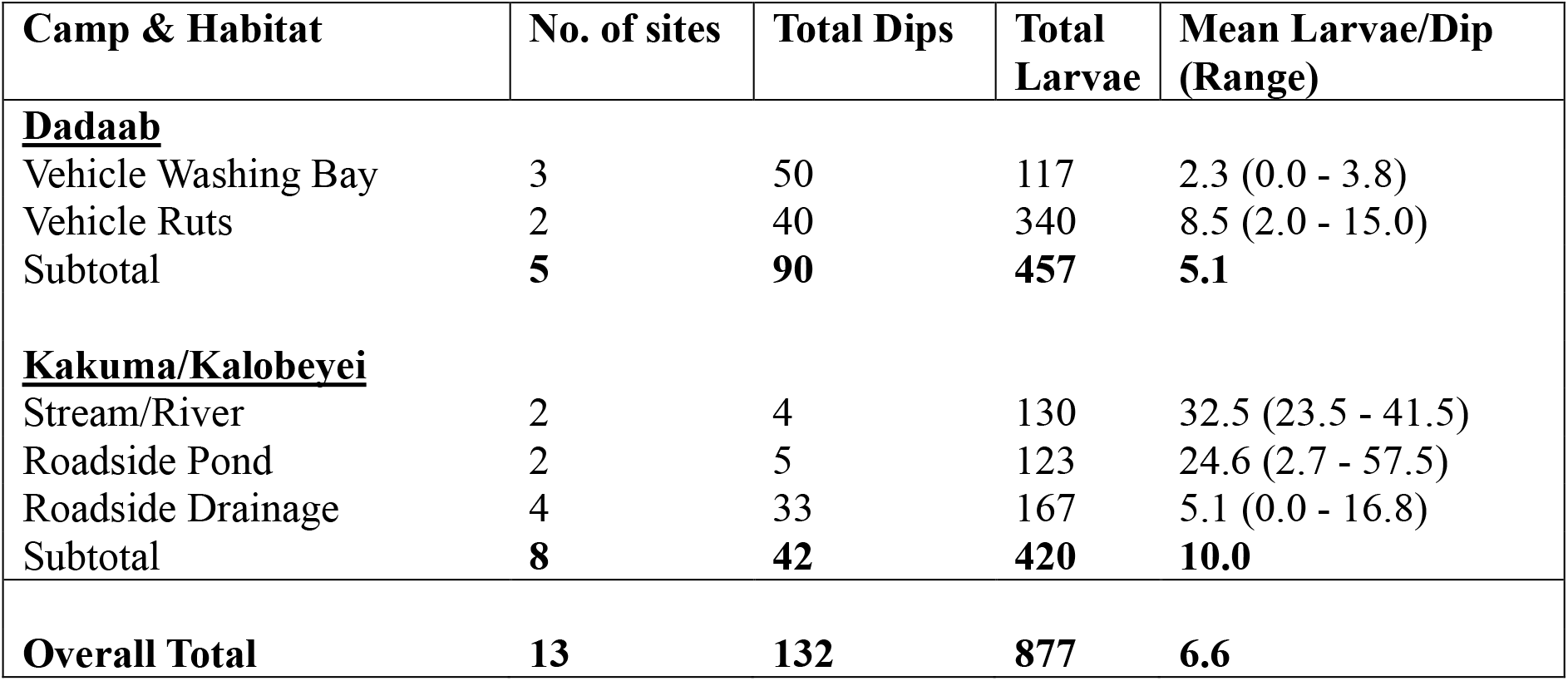
Summary of Anopheles larval habitat type in refugee camps in Kenya.

In Dadaab, larval production was confined to anthropogenic habitats. Vehicle washing bays and water-filled vehicle ruts were the primary larval sources, yielding a sub-total of 457 larvae. While vehicle ruts were the most prolific habitat type in this camp (mean density: 8.5 larvae/dip), the overall mean density for Dadaab was 5.1 larvae/dip (Table 1).

In contrast, breeding sites in Kakuma and Kalobeyei were more varied and productive, comprising natural and peri-domestic habitats (Figure 1B&C). These sites yielded a higher overall mean density of 10.0 larvae/dip. As detailed in Table 1, the most productive habitat categories were streams/riverbeds (mean density: 32.5 larvae/dip) and roadside ponds (mean density: 24.6 larvae/dip). The single most productive habitat identified in the entire study was a roadside pond in Kakuma, which reached a density of 57.5 larvae/dip. The highly variable productivity of roadside drainages, which ranged from zero to 16.8 larvae/dip, underscores the heterogeneous distribution of breeding sites within the camp environment.

From 877 larvae collected, DNA was extracted successfully from 728 and subjected to molecular genotyping for *An. gambiae, An. funestus* species complexes and *An. stephensi* and ITS-2 amplicon sequencing. Notably, no specimens of the *Anopheles funestus* group or *Anopheles stephensi* were identified in any of the camps. Overall, Anopheles arabiensis dorminated larval collection in refugee camps(59%) followed by *An. coluzzii* (35%), An rufipes (1%) and lastly a single specimen of *An. gambiae s*.*s*. identified. Approximately 5% of mosquitoes collected could not be identified by combination of PCR based methods nor ITS-2 sequencing using the sanger sequencing approach (Figure 2A, Table S1 and Table S2).

The vectors identified were spatially distributed as follows. In Dadaab, *An. arabiensis* was the predominant species, accounting for 94% (n = 352) of the collections, with only a single specimen of *An. gambiae* s.s. identified. In Kakuma, *An. coluzzii*. dominated (84%, n = 222), while *An. arabiensis* represented 10% (n = 27). In Kalobeyei, *An. arabiensis* (54%, n=47) and *An. coluzzii* (34%, n=30) and *An. rufipes* (7%, n=6) were the primary species (Figure 2B). In addition, samples from western Kenya (Luyeshe and Maseno) were included for comparative purposes and only identified *An. gambiae s*.*s*. (Figure S1).

**Figure 2.**
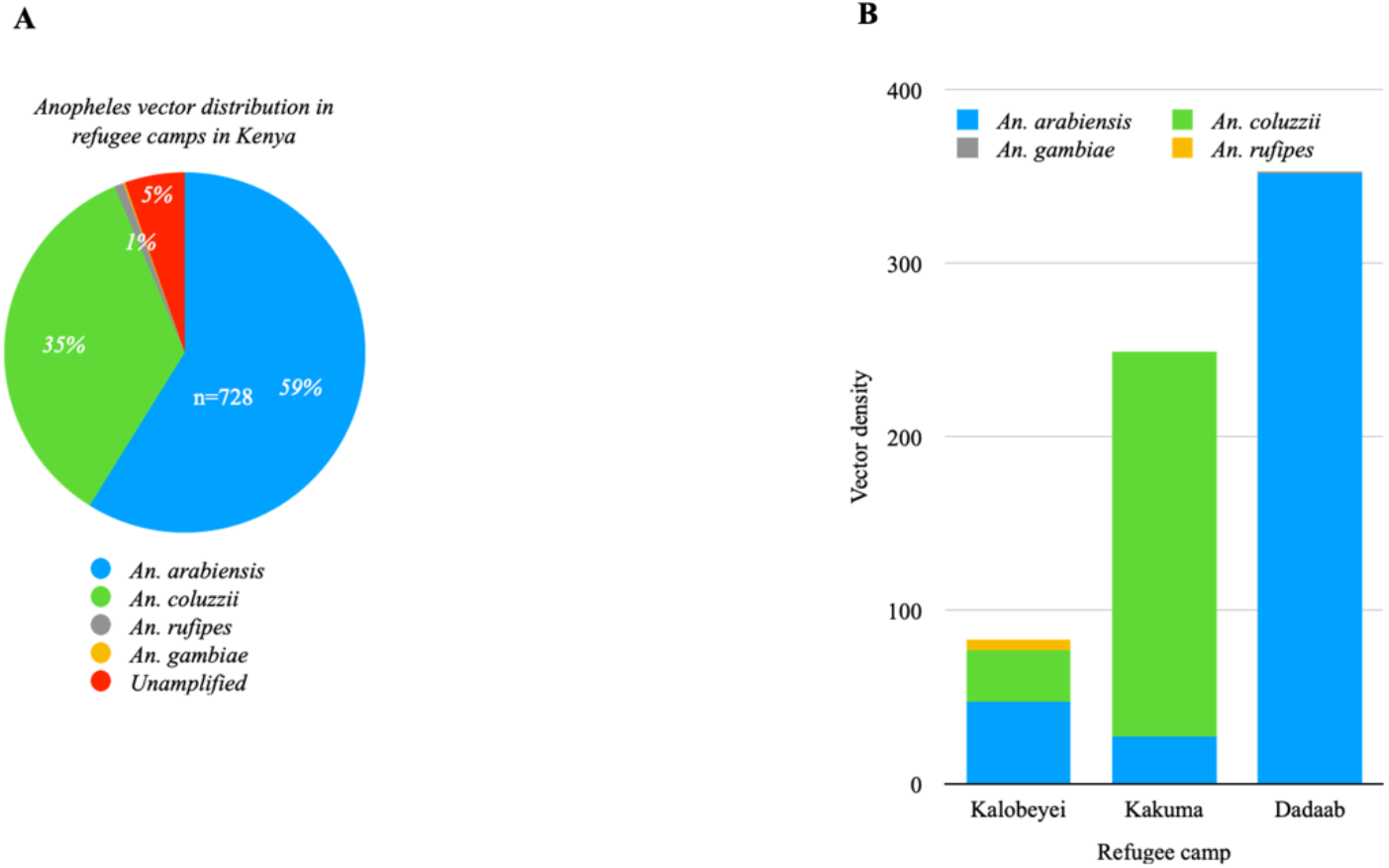
Species composition of Anopheles mosquitoes in Kakuma, Kalobeyei, and Dadaab refugee camps, Kenya. (A) Overall relative abundance of Anopheles species identified by PCR and ITS2 sequencing. (B) Spatial distribution and abundance of the predominant species, *An. coluzzii* and *An. arabiensis*, across individual sampling sites.

### Frequency of *kdr* mutations in *vgsc* of *An. gambiae s*.*l*

The *kdr*-L1014F mutation was detected at high frequencies in *An. coluzzii* from Kakuma and Kalobeyei, with allelic frequencies of 50% and 63%, respectively. In contrast, *An. arabiensis* exhibited lower frequencies of the L1014F mutation (10% and 30% in Kakuma and Kalobeyei, respectively). Neither the L1014S nor L1014F mutation was observed in specimens from Dadaab. High levels of insecticide resistance mutations were thus limited to *An. coluzzii*. in Turkana County and absent in Garissa (Table 2).

**Table 2.** *kdr-*West (L1014F) genotype distribution and allelic frequencies in Anopheles populations across study sites.

| Site | *kdr*-West(L1014F) |  |  |  |  |  |  |
| --- | --- | --- | --- | --- | --- | --- | --- |
|  | Species | Genotype count |  |  |  | Allelic frequency |  |
|  |  | # | RR | RS | SS | R | S |
| Dadaab | *An. arabiensis* | 322 | 0 | 0 | 322 | 0 | 1 |
| kakuma | *An. coluzzii* | 58 | 3 | 52 | 3 | 0.5 | 0.5 |
| Kakuma | *An. arabiensis* | 26 | 5 | 4 | 17 | 0.27 | 0.73 |
| Kalobeyei | *An. coluzzii* | 27 | 9 | 16 | 2 | 0.63 | 0.37 |
| Kalobeyei | *An. arabiensis* | 40 | 0 | 8 | 32 | 0.1 | 0.9 |

## Discussion

Malaria outbreaks in Turkana County have increased in recent years, with year-round *P. falciparum* transmission that is not strongly correlated with rainfall patterns[28]. Additionally, *P. vivax* infections have been reported among semi-nomadic populations in Turkana and northeastern Kenya[29] who have been implicated in facilitating parasite spread through their migration routes. Shifts in mosquito vector populations are known to influence malaria transmission [30, 31]. Understanding bionomics of vector populations at the local scale is a critical first step for elucidating transmission dynamics and informing targeted intervention. This study investigated the larval ecology of malaria vectors in Kakuma and Dadaab refugee camps, revealing significant changes in vector composition. In Kakuma this represents a significant shift from the previously documented vector population from 2005-2006, which was reported to be homogeneously *An. arabiensis* [26]. The population has now changed to a more heterogeneous population comprising *An. coluzzii, An. arabiensis* and *An rufipes* two decades later. Notably, *An. rufipes* was identified through the ITS2 sequencing approach after being missed by standard PCR, underscoring the value of complementary molecular methods for accurate species resolution. The role of this vector in the Kakuma-Kalobeyei complex cannot be overlooked, as it may contribute to the local transmission dynamics. While malaria vector populations shifts have been documented in various settings, they have tended towards a decline in *An. gambiae s*.*s* and an increase in *An. funestus* as the dominant species[32]. The absence of *An. funestus* and *An. stephensi* in our larval collections, despite their known presence in other parts of Kenya, suggests that the specific breeding habitats in these camps— dominated by temporary sunlit pools and vehicle ruts may not be suitable for these species, which prefer more permanent, vegetated, or container-based habitats respectively. These ecological shifts are largely attributed to widespread insecticide use[32]. Despite year-round malaria transmission, vector surveillance In Kakuma has been limited, leaving the primary malaria vectors poorly characterized. Concurrently, the camp’s human population has surged over the past two decades, prompting establishment of the Kalobeyei settlement.

These anthropogenic changes, along with environmental and ecological factors may have created conditions conducive for proliferation of *An. coluzzii* [33]. It is theorized that the speciation of *An. gambiae* and *An. coluzzii* is driven not only by genetics but ecological factors including quality of aquatic habitats[19]. The later emergence of *An. coluzzii* is associated with the diversification of aquatic habitats including those of marginal quality, for which it appears predisposed[19]. This is to some extent reflected in the diversity and unusual habitats within which Anophelines were found breeding in Kakuma. With regards to infection, susceptibility to *Plasmodium falciparum* of all three vector species varies depending on local environment, genetic factors and parasite strain. However, high *P. falciparum* infection rates have been reported in *An. coluzzii* and in one study *P. vivax* infection was documented [34]. In Kakuma, the current vector composition represents a notable change from the previously documented population from 2005-2006, which was reported to be homogeneously *An. arabiensis* [26]. The population now comprises a more heterogeneous mixture of *An. coluzzii, An. arabiensis*, and *An. rufipes*. Given the lack of continuous longitudinal data, this change could represent either a local population shift or the recent introduction and subsequent establishment of *An. coluzzii* in the region. The presence and establishment or expansion of *An. coluzzii* in the region appears to be a relatively recent phenomenon, as it was not detected during vector surveillance in Kakuma fifteen years ago [26]. Therefore, its role in past and current malaria outbreaks in Turkana County is still poorly understood warranting further investigation, particularly given conflicting evidence on its vectorial capacity and susceptibility to *Plasmodium* infections[35, 36].

In contrast, *An. arabiensis* remained the dominant vector in Dadaab. Unlike Kakuma, Dadaab experiences little to no malaria transmission. The underlying mechanisms for this low transmission, despite the presence of a competent vector, are not fully understood. Potential contributing factors could include the zoophilic tendencies of local *An. arabiensis* populations, the influence of endosymbiotic microorganisms, or other genetic, environmental, and anthropogenic factors that may reduce transmission efficiency[37, 38]. However, the specific drivers in this context remain unclear, underscoring the need for dedicated studies to elucidate the malaria transmission dynamics and the relative contribution of different factors in Dadaab Overall, the identity of approximately 5% of the samples could not be resolved using standard *An. gambiae s*.*l*. and *An. funestus s*.*l*. species complex, An stephensi species specific PCR or ITS2 amplicon sequencing identification protocols. This includes definitively ruling out *An. stephensi*, for which we employed a specific PCR assay[14], confirming it was not present among these unresolved specimens. The fact that *An. rufipes* was only detected through sequencing highlights a key limitation of standard PCR assays and highlights the critical need to employ complementary sequencing and genomic tools to fully characterize anopheline diversity. This approach is essential to uncover potentially overlooked secondary vectors whose role in transmission may currently be underestimated.

Regarding insecticide resistance, high frequencies of the *kdr*-L1014F mutation were observed in *An. coluzzii* from Kakuma and Kalobeyei suggesting a move towards fixation, with most mosquitoes exhibiting either homozygous or heterozygous genotypes. Lower frequencies of the mutation were detected in *An. arabiensis* from the same sites, while no kdr mutations were found in specimens from Dadaab. These findings suggest selection pressure favoring insecticide resistance genotypes in Turkana County, potentially driven by local insecticide use or the introduction of resistant mosquitoes from neighboring regions, such as eastern Uganda, where high levels of resistance have been documented[39]. However, this study focused solely on genotypic resistance; phenotypic assays are needed to confirm the susceptibility of these populations to insecticides used in vector control programs.

Long-lasting insecticidal nets (LLINs) and indoor residual spraying (IRS) remain critical for malaria control in refugee camps, but their efficacy depends on mosquito susceptibility to insecticides. The emergence of pyrethroid resistance in Turkana County, evidenced by high *kdr*-L1014F frequencies, demands integrated vector management strategies and continuous monitoring of *kdr* allele frequencies to mitigate this threat.

While our study provides key insights into larval ecology across three refugee settlements, we acknowledge limitations inherent to larval-based surveillance. Larval collections are subject to sampling bias, as cryptic breeding sites, particularly those of *Anopheles funestus* and secondary vectors, often evade detection despite confirmed adult presence. Consequently, our data reflect larval diversity within *accessible* habitats rather than exhaustive vector composition. Nevertheless, in Turkana (Kakuma/Kalobeyei) and Dadaab, where anthropogenic sites (water storage containers, drainage ditches) sustain dominant *An. coluzzii* and *An. arabiensis* populations, larval source management (LSM) emerges as a high-priority intervention. Targeted LSM (e.g., container covering, habitat drainage, site-specific larviciding) could effectively suppress these container-breeding *gambiae*-complex vectors. In contrast, settings with confirmed *funestus*-group dominance would require alternative approaches due to their association with cryptic, vegetated aquatic habitats. Thus, our findings strongly support LSM in semi-arid humanitarian contexts where *gambiae*-complex vectors dominate identifiable artificial habitats, while underscoring the need for complementary adult surveillance to resolve cryptic vector dynamics.

## Conclusion

This study documents significant ecological shifts in malaria vectors across Kenya’s arid-zone refugee settlements, revealing the emergence of *Anopheles coluzzii* alongside *An. arabiensis* in Turkana County, linked to anthropogenic habitat modifications that enable year-round transmission—while Dadaab exhibits persistent *An. arabiensis* dominance without proportional malaria burden, suggesting unexplained local refractoriness. Critically, high *kdr*-L1014F frequencies in Turkana’s *An. coluzzii* (upto 63%) pose an immediate threat to pyrethroid-based interventions (LLINs/IRS), necessitating urgent resistance management through next-generation tools and phenotypic validation. Furthermore, the unresolved identity of 5% of specimens underscores the need for enhanced surveillance using whole-genome sequencing or

Matrix-Assisted Laser Desorption/Ionization Time-of-Flight (MALDI-TOF). These findings collectively highlight the urgency for habitat-focused vector control in semi-arid humanitarian contexts: where *gambiae*-complex vectors dominate identifiable artificial breeding sites (water storage containers, drainage ditches), targeted larval source management (container covering, site-specific larviciding) offers a critical strategy to disrupt transmission at its source, complementing insecticide resistance countermeasures to protect crisis-affected populations.

## Acknowledgement

We thank the technical and field staff: Festus Yaa, Gabriel Nzai, and Julius Tineja, who helped with mosquito sample collection in the field.

## Author contributions

M.K.R. conceptualisation, methodology, formal analysis, funding acquisition, writing— original draft. M.M. conceptualisation, methodology, reviewing and editing. C.W. methodology, reviewing and editing. L.I.O. funding acquisition and reviewing and editing. J.M. reviewing and editing. R.W.S. reviewing and editing. B.B., K.G., A.D., M.T. and A.O. sample collection and processing, laboratory analysis and reviewing and editing. All authors read and approved the manuscript.

## Ethics approval and consent to participate

The study was approved by the KEMRI Scientific and Ethics Review Unit (SERU) with the protocol number: 337 KEMRI/SERU/CGMR-C/024/3148.

## Consent for publication

This manuscript is published with the permission of the Director-General of the Kenya Medical Research Institute.The funding bodies had no role in the design, data collection, and drafting of the manuscript.

## Availability of data and materials

Most of the dataset used for analysis is available in the manuscript. We withheld the geo-data that may predispose individual homesteads to a high risk of identifiability. However, they are under the custodianship of the KEMRI Wellcome Trust Data Governance Committee and are accessible upon request addressed to that committee.

